# The meninges enhance leukemia survival in cerebral spinal fluid

**DOI:** 10.1101/750513

**Authors:** Patrick Basile, Leslie M. Jonart, Maryam Ebadi, Kimberly Johnson, Morgan Kerfeld, Peter M. Gordon

**Author notes:** Contact information for corresponding author: Peter Gordon, M.D./Ph.D. Division of Pediatric Hematology/Oncology University of Minnesota 420 Delaware St SE, MMC 366 Minneapolis, MN, 55455.

## Abstract

Central nervous system (CNS) relapse is a common cause of treatment failure in patients with acute lymphoblastic leukemia (ALL) despite current CNS-directed therapies that are also associated with significant short and long-term toxicities. Herein, we showed that leukemia cells exhibit decreased proliferation, elevated reactive oxygen species (ROS), and increased cell death in CSF both *in vitro* and *in vivo*. However, interactions between leukemia and meningeal cells mitigated these adverse effects. This work expands our understanding of the pathophysiology of CNS leukemia and suggests novel therapeutic approaches for more effectively targeting leukemia cells in the CNS.

---

Acute lymphoblastic leukemia (ALL) is the most common pediatric cancer. While significant progress has been made in the therapy of leukemia, several obstacles still render cure either harder to obtain or to come at a higher cost. Even in the setting of adequate upfront risk stratified therapy, central nervous system (CNS) relapse is a common cause of treatment failure and current CNS-directed therapies are associated with significant morbidities^1^. However, a better understanding of how the CNS microenvironment influences leukemia biology may lead to more effective and less toxic therapies for CNS leukemia^2,3^. A unique characteristic of the CNS microenvironment is that leukemia cells are bathed in cerebral spinal fluid (CSF). Despite being derived from plasma, CSF exhibits significant differences in its electrolyte and protein composition, including over two orders of magnitude less proteins than in plasma^4,5^.

To investigate the effect of CSF on leukemia cell biology, we began by assessing the effect of CSF on leukemia cell proliferation and viability. Two different leukemia cell lines (Jurkat, NALM-6), representing both a T- and B-cell leukemia, were cultured in either standard growth media (RPMI supplemented with 10% fetal bovine serum (FBS)) or human cerebrospinal fluid. Leukemia cell proliferation was assessed daily by both manual cell counts and with the CellTiter-Glo Viability Assay (Promega). Both measures showed that leukemia cell proliferation was significantly diminished in CSF relative to standard RPMI tissue culture media (Figure 1A-B). We then used a fixable viability dye and flow cytometry to compare leukemia cell death in RPMI and CSF. As shown in Figure 1C, both leukemia cell lines showed a significant decrease in viability when cultured in CSF (Figure 1C). We then used BH3 profiling, a functional apoptosis assay that predicts the propensity of a cell to undergo apoptosis, to further characterize the effect of CSF on leukemia cells^6^. As shown in Figure 1D, leukemia cells cultured in CSF retained less cytochrome C after treatment with the pro-apoptotic BIM peptide than leukemia cells in media. This result is consistent with leukemia cells in CSF being more poised to undergo apoptosis. We next asked whether the increased leukemia cell death in CSF was secondary to a toxic component in the CSF or if there is a required nutrient or substrate missing in CSF compared to media. To address this question, we cultured NALM-6 and Jurkat leukemia cells in RPMI media, regular CSF, or CSF supplemented with either RPMI base powder or 10% FBS. We then measured viability daily using the CellTiter-Glo Viability assay. As shown in Figure 1E, CSF supplemented with FBS, but not RPMI powder, showed slightly improved viability compared to baseline CSF, but not to the levels of RPMI media. These results suggest that both the presence of a component within CSF and the lack of a substrate within CSF, supplied by FBS, contribute to the poor proliferation and survival of leukemia cells in CSF. Given the altered level of redox proteins in the CSF^7^, we hypothesized that ROS levels may be altered in leukemia cells in CSF. Moreover, while ROS may have leukemogenic effects, excessive ROS and oxidative stress are associated with priming mitochondrial apoptosis and this excess stress is the mechanism of action of some chemotherapeutics, including those used or tested in the therapy of ALL^8–10^. We used the CellRox Green stain (Life Technologies) with flow cytometry to measure ROS levels in viable Jurkat and NALM-6 leukemia cells cultured in either RPMI or CSF. The median fluorescence intensity (MFI) of ROS increased in both leukemia cell lines in CSF whereas it remained stable in RPMI (Figure 1F-G). We then measured mitochondrial ROS using the MitoSox stain (Thermo Fisher, Waltham, MA USA) in order to try and localize the sub-cellular origin of the ROS in leukemia cells in CSF. As shown in Figure 1H, Jurkat, but not NALM-6, leukemia cells showed increased mitochondrial ROS, suggesting that mitochondria may be at least one sub-cellular source for the elevated ROS observed in leukemia cells in CSF.

**Figure 1.**
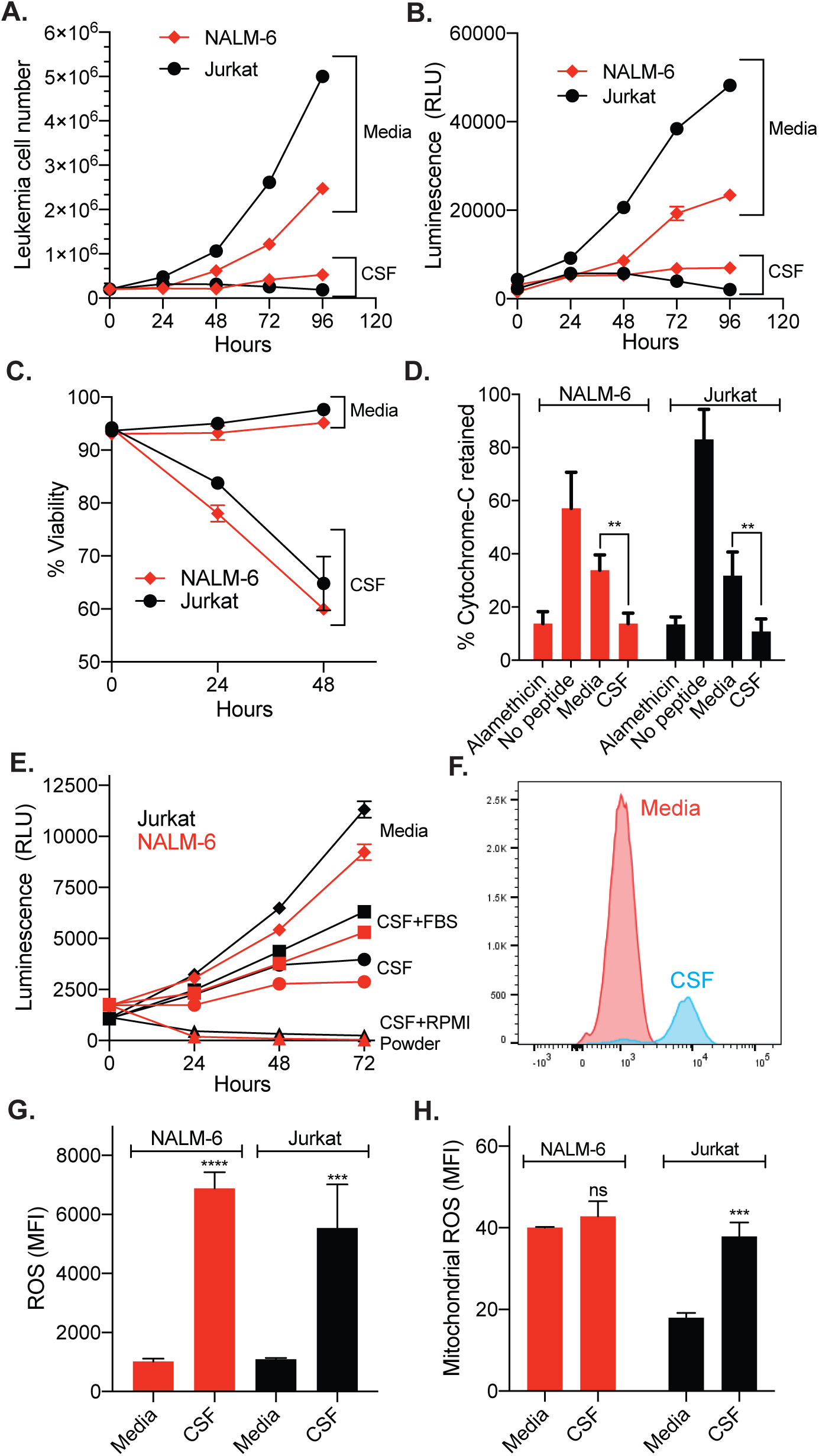
Leukemia cells in CSF exhibit decreased proliferation and viability but increased ROS. (A-C) NALM-6 and Jurkat leukemia cells were cultured in either regular media (RPMI and FBS 10%) or human CSF. The effect of CSF on leukemia cells was then assessed daily using manual cell counts and a hemocytometer (A), the CellTiter-Glo Viability Assay and a microplate reader capable of measuring luminescence (B), and a fixable viability dye and flow cytometry (C). (D) BH3 profiling was performed with the BIM peptide on NALM-6 and Jurkat leukemia cells cultured in regular media or human CSF for 48 hours. Cytochrome c retention was measured by flow cytometry. Alamethicin is a peptide antibiotic that permeabilizes the mitochondria membrane and is a positive control. (E) NALM-6 and Jurkat leukemia cells were cultured in either regular media (RPMI and FBS 10%), human CSF, or human CSF supplemented with either FBS 10% or RPMI powder. Leukemia cell proliferation was then measured daily with the CellTiter-Glo Viability Assay. (F-H). NALM-6 and Jurkat leukemia cells were cultured in either regular media (RPMI and FBS 10%) or human CSF. After 72 hours, leukemia cells were stained with either CellROX green (F-G) or MitoSox red dye (H) and then median fluorescent intensity (MFI) assessed by flow cytometry. A representative flow cytometry plot for Jurkat leukemia cells stained with CellROX dye is shown in (F). For all graphs, data are the mean +/− SD from three independent experiments and *P*: **, <0.01, ****, <0.0001 by ANOVA or t-test. *ns*, not significant.

Together these data suggest that leukemia cell proliferation and survival are poorly supported by the liquid milieu of the CNS microenvironment. Accordingly, we then asked whether direct interactions between leukemia cells and meningeal cells could mitigate this diminished ability of CSF to support leukemia cell survival. We selected the meninges for these experiments because histopathological examinations from leukemia patients and mice transplanted with human leukemia cells have identified the meninges as a primary site of leukemia involvement in the CNS^11,12^. Leukemia cell lines were grown suspension or adherent to primary, human meningeal cells in the presence of CSF. After 72 hours, we assessed leukemia viability and ROS using a fixable viability dye and CellRox stain, respectively. As shown in Figure 2A-B, adhesion to meningeal cells in CSF significantly increased leukemia cell viability and attenuated ROS levels relative to the same leukemia cells grown in suspension in CSF. Similar to the results with leukemia cell lines, meningeal cells also significantly enhanced the survival of primary pre-B leukemia cells that had never been cultured *ex vivo* (Figure 2C). Further supporting the importance of direct cell-cell interactions, adhesion of leukemia cells to tissue culture plates treated with recombinant fibronectin (RetroNectin) did not enhance leukemia cell survival in CSF (Supplemental Figure 1). We next used a leukemia xenotransplantation system to assess the *in vivo* effect of CSF on leukemia viability. Immunocompromised (NSG) mice were transplanted with NALM-6 leukemia cells. After ~3 weeks and systemic leukemia development, mice were euthanized and CSF, bone marrow, meninges and blood harvested. Assessment of leukemia cell viability in each tissue showed a marked reduction in leukemia cell viability in CSF compared to the other three tissues (Figure 2D). Together these data suggest that interactions between leukemia and meningeal cells enhance leukemia survival in CSF. Supporting the potential clinical translation of these results, we also found that cell adhesion inhibitors AMD3100 (CXCR4 antagonist) and A205804 (ICAM-1 and E-selectin inhibitor) disrupted leukemia-meningeal adhesion in co-culture and that the displaced leukemia cells exhibited decreased viability in CSF (Supplemental Figure 2).

**Figure 2.**
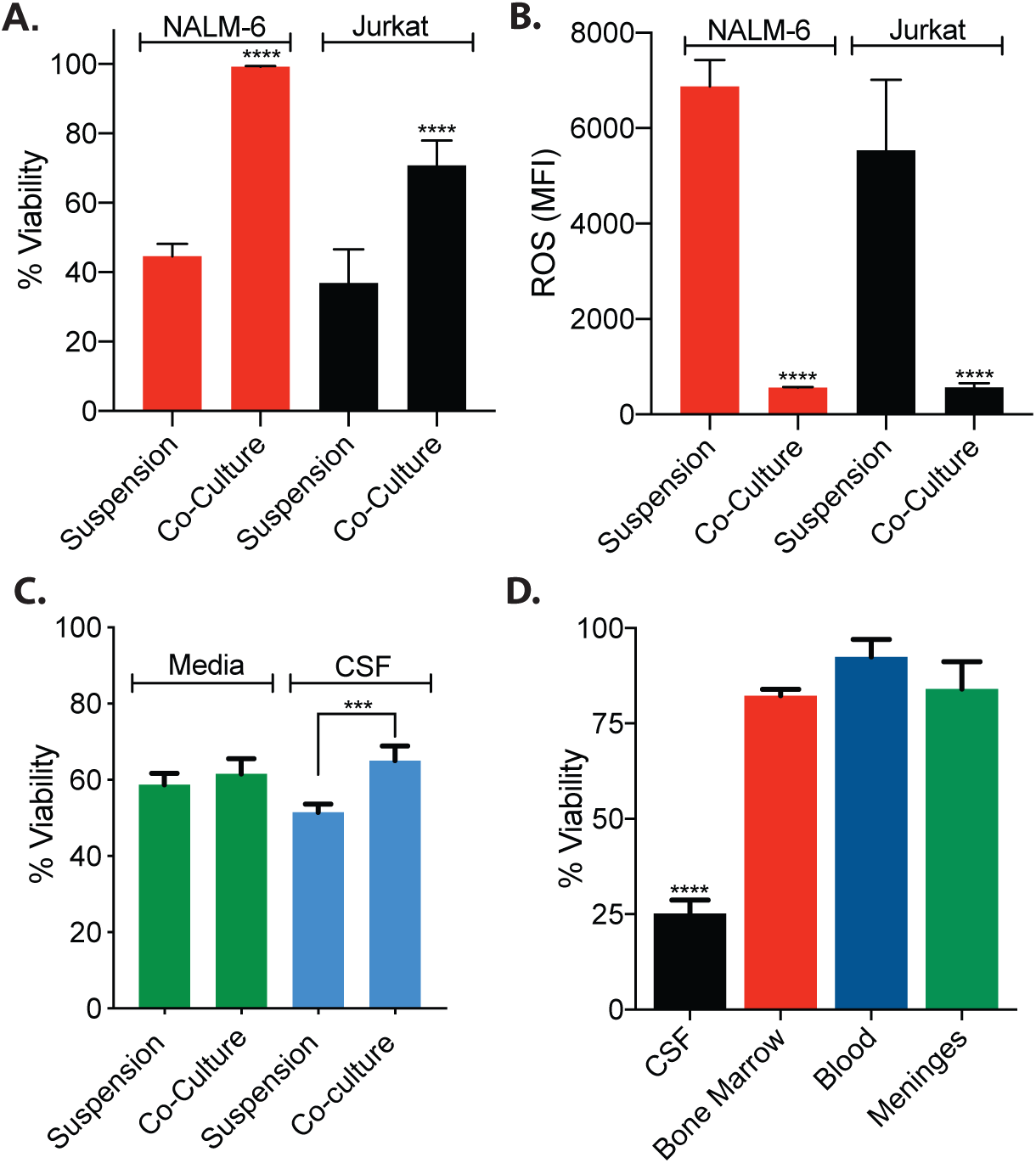
The meninges enhance leukemia viability both in vitro and in vivo. (A-C) NALM-6, Jurkat, and primary pre-B (C) leukemia cells were cultured in human CSF either in suspension or adherent to primary, human meningeal cells. After 48 (C) or 72 hours (A-B), leukemia cell viability (A, C) and ROS levels (B) were assessed using a fixable viability dye or CellROX green dye, respectively, and flow cytometry. Data are the mean +/− SD from three experiments and *P*: ***, <0.001 and ****, <0.0001 by ANOVA or t-test. (D) NSG mice (N=3) were transplanted with NALM-6 leukemia cells (2×10^6^ cells). After ~3 weeks, mice were euthanized and CSF, bone marrow, meninges, and peripheral blood harvested. Leukemia cell viability in each tissue or fluid was then assessed using a fixable viability dye and flow cytometry. *P*: ****, <0.0001 by t-test.

An alternative explanation for the ability of leukemia cells to persist in the CNS despite the deleterious effects of CSF is that rare leukemia cells may exhibit distinct properties that enhance survival and proliferation in the CSF and that these cells are subsequently selected for within the CNS microenvironment. Arguing against this possibility, sequencing of paired B-cell leukemia samples from the bone marrow and CSF of individual patients or xenotransplanted mice showed similar clonal architecture patterns between the two different microenvironments^11,13^. This suggests that most B-cell leukemia clones are capable of trafficking to and persisting within the CNS, although phenotypic differences or selection within clones could contribute to leukemia persistence in CSF^14^. However, despite >3 weeks of continuous *ex vivo* culture we were unable to select for the emergence of leukemia cells with enhanced proliferation and survival in CSF (data not shown).

In summary, we have shown that interactions between leukemia and meningeal cells attenuates the deleterious effects of CSF on leukemia proliferation and survival. This work provides new insights into CNS leukemia pathophysiology and illustrates the unique effects different niches, and components of a niche, exert on leukemia biology. These results also suggest that the laboratory examinations of CSF for leukemia cells should be performed in a timely manner as any delays could diminish the likelihood for detecting viable leukemia cells and, thus, alter leukemia staging or therapy response assessments. Finally, the ability of cell adhesion inhibitors to disrupt leukemia-meningeal adhesion with resulting leukemia cell death in CSF suggests it may be possible to leverage these results into new therapies for CNS leukemia. Further supporting this approach, drugs that disrupt interactions between leukemia cells and components of the bone marrow have shown pre-clinical efficacy in sensitizing leukemia cells to chemotherapy and are undergoing clinical testing, development, and optimization^15^. Moreover, niche disruption may be more efficacious in the CNS than in the bone marrow because of the less supportive environment of the CSF relative to the blood or serum.

## Acknowledgements

This work was supported in part by the Children’s Cancer Research Fund (PMG), the Timothy O’Connell Foundation (PMG), and an American Cancer Society Institutional Research Grant (PMG). PB was partially supported by NIH Training Grant T32 CA099936.

## Contributions

P.B. designed experiments, analyzed data, and prepared figures. P.B., L.J., M.K. and K.J. performed experiments and analyzed data. P.M.G. designed the study, oversaw the laboratory investigations, and wrote the manuscript. All authors reviewed, edited and approved the final version of the manuscript. We thank Leah Kann for assisting with CSF procurement.

