## Supplemental Materials for "The meninges enhance leukemia survival in cerebral spinal fluid"

**Supplementary Information**

***Materials and Methods***

***Reagents and Cell Culture:*** Jurkat (T-cell) and NALM-6 (B-cell) leukemia cell lines were obtained from American Type Culture Collection (ATCC) and DSMZ, respectively. Primary pre-B leukemia cells (Sample CBAB-62871-V1) were obtained from the Public Repository of Xenografts (PRoXe[1]). The leukemia cell lines were cultured in RPMI media supplemented with Fetal Bovine Serum (FBS; Seradigm) 10% and Penicillin-Streptomycin. Primary meningeal cells were obtained from ScienCell and cultured in meningeal media supplemented with FBS 2%, growth supplement, and Penicillin-Streptomycin. Human cerebral spinal fluid (CSF), pooled from healthy donors, was obtained from Lee Biosolutions. Additional CSF was obtained from non-infectious patients undergoing neurosurgical procedures at the University of Minnesota Masonic Children's Hospital after approval from the University of Minnesota IRB (IRB ID#00001210). CSF was filtered and centrifuged to remove any cellular components or debris prior to use. RPMI powder was obtained from ThermoFisher and dissolved in CSF according to manufacturer's instructions. Tissue culture plates were treated with RetroNectin (Clonetch, recombinant fibronectin fragment) according the manufacturer's instructions. AMD3100 (CXCR4 antagonist) and A205804 (ICAM-1 and E-selectin inhibitor) were obtained from Sigma-Aldrich.

***Proliferation, viability and ROS assays:*** In 96-well plate format, leukemia cells were treated as described for 48 hours and then viability assessed with the CellTiter-Glo Luminescent Cell Viability Assay (Promega; Madison, WI) and a Molecular Devices SpectraMax plate reader. All experiments were performed in triplicate with at least 3 wells per condition. Manual cell counts were performed using trypan blue staining and a hemocytometer. To assess viability, leukemia cells were treated as described, stained with fixable viability dye (ThermoFisher), and analyzed by flow cytometry. ROS was measured by staining leukemia cells with either CellROX green reagent (ThermoFisher) or MitoSox Red reagent (ThermoFisher) and assessing with flow cytometry[2].

***Leukemia co-culture:*** 2.5 x $10^5$ leukemia cells were labeled using Cell Trace Violet (Invitrogen) to enable identification of leukemia cells. The stained cells were then plated onto 60% confluent primary meningeal cells (24 well plate) and grown in CSF for 48-72 hours. The CSF was then aspirated and the co-cultures were gently washed with ice cold PBS. Cells were then trypsinized with 0.05% Trypsin and manual pipetting prior to staining for viability and ROS as previously described. For experiments with cell adhesion inhibitors, leukemia and meningeal cells were co-cultured in CSF with no drug, AMD3100 100 μM or A205804 1 μM. After 24 hours, the non-adherent leukemia cells were transferred to another well in the absence of meningeal cells and additional CSF was added to the co-culture wells. After an additional 24 hours, leukemia cells adherent to the meningeal cells in co-culture wells were removed by manual pipetting and counted by flow cytometry and Count Bright Absolute counting beads (ThermoFisher). The viability of non-adherent leukemia cells in CSF was measured using annexin-V/7-AAD staining and flow cytometry.

**BH3 Profiling:** BH3 profiling was performed as described[3,4]. Leukemia cells were grown in regular media or CSF for 48 hours and then suspended in membrane extraction buffer containing digitonin 100 μg/mL (Sigma-Aldrich). The leukemia cells were then incubated with the BIM peptide 50 μM (GenScript) for 90 minutes. After incubation, leukemia cells were fixed with paraformaldehyde 4%, permeabilized, stained for cytochrome c (eBioscience), and analyzed by flow cytometry.

***Xenograft studies:*** NSG (NOD.Cg-*Prkdcscid, Il2rgtm1Wjl*/SzJ; Jackson Labs, Bar Harbor, ME) mice were housed under aseptic conditions and received autoclaved cages, bedding material, water, bottles, and irradiated food. Mouse care and experiments were in accordance with protocols approved by the Institutional Animal Care and Use Committee at the University of Minnesota. 4-7 week old mice were injected intravenously via the tail vein with ~$2x10^6$ leukemia cells. After systemic leukemia development (~3 weeks), peripheral blood was collected via facial vein bleeding and then red cell lysed using RBC Lysis Buffer (eBioscience). After euthanasia the mice were cardiac perfused with PBS and CSF was obtained from mice as described[5]. Briefly, mice were euthanized and a scalp incision was made at the midline to expose the dura mater overlying the cisterna magna. Under a dissection microscope, a tapered, pulled glass capillary tube was then inserted through the dura and into the cisterna magna to obtain clear CSF. The meninges were then removed using a dissecting microscope, and dissociated by gently washing through a 0.40 µm filter (Millipore). Finally, the femurs were removed, crushed with mortar and pestle, and red cell lysed with RBC Lysis Buffer. Cells were then stained with fluorescent antibodies against CD19 (NALM-6; eBioscience) or CD3 (Jurkat, eBioscience), a fixable viability dye (ThermoFisher), and analyzed by flow cytometry using a BD FACSCanto or BDFACSCalibur.

***Statistical analysis:*** Results are shown as the mean plus or minus the SD of the results of at least 3 replicates. The Student's *t*-test or ANOVA were used for statistical comparisons between groups and were calculated using GraphPad Prism 8 software (GraphPad Software, La Jolla, CA). *P*-values less than 0.05 were considered statistically significant

***Supplemental Figure Legends***

***Supplemental Figure 1: Adhesion to fibronectin does not enhance leukemia cell survival in CSF.*** NALM-6 and Jurkat leukemia cells were cultured in CSF either adherent to tissue culture plates that were pre-treated with recombinant human fibronectin fragment (RetroNectin) or in suspension in non-treated plates. After 48 hours, leukemia proliferation was measured with the CellTiter-Glo Viability Assay and a microplate reader. *P*: *, <00.5; **, <0.01 by ANOVA.

***Supplemental Figure 2: Disrupting leukemia-meningeal adhesion in CSF increases leukemia cell death.*** *(*A-B) NALM-6 leukemia cells were co-cultured with meningeal cells in CSF with AMD3100 100 μM (CXCR4 antagonist), A205804 1 μM (ICAM-1 and E-selectin inhibitor), or no drug control. After 24 hours of drug treatment, non-adherent leukemia cells were removed and quantitated (A). NALM-6 viability was then measured with annexin-V/7-AAD staining and flow cytometry after an additional 24 hours in CSF in suspension (B). *P*: *, <00.5; **, <0.01; ****, <0.0001 by ANOVA.

***References***


1. Townsend EC, Murakami MA, Christodoulou A, Christie AL, Köster J, DeSouza TA, et al. The Public Repository of Xenografts Enables Discovery and Randomized Phase II-like Trials in Mice. Cancer Cell. 2016 Apr 11;29(4):574–86.
2. Kauffman M, Kauffman M, Traore K, Zhu H, Trush M, Jia Z, et al. MitoSOX-Based Flow Cytometry for Detecting Mitochondrial ROS. ROS. 2016;
3. Ryan J, Montero J, Rocco J, Letai A. iBH3: simple, fixable BH3 profiling to determine apoptotic priming in primary tissue by flow cytometry. Biol Chem. 2016 Jul 1;397(7):671–8.
4. Ryan J, Letai A. BH3 profiling in whole cells by fluorimeter or FACS. Methods. 2013 Jun 1;61(2):156–64.
5. Liu L, Duff K. A technique for serial collection of cerebrospinal fluid from the cisterna magna in mouse. J Vis Exp. 2008 Nov 10;(21).

### Supplemental Figure 1

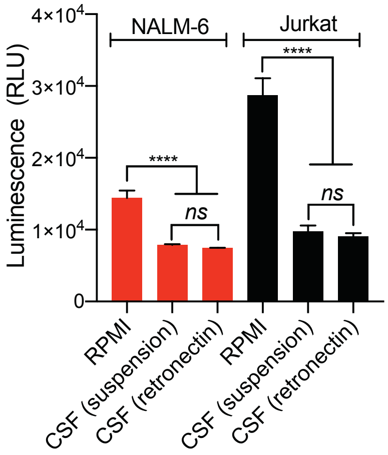

**Supplemental Figure 2**

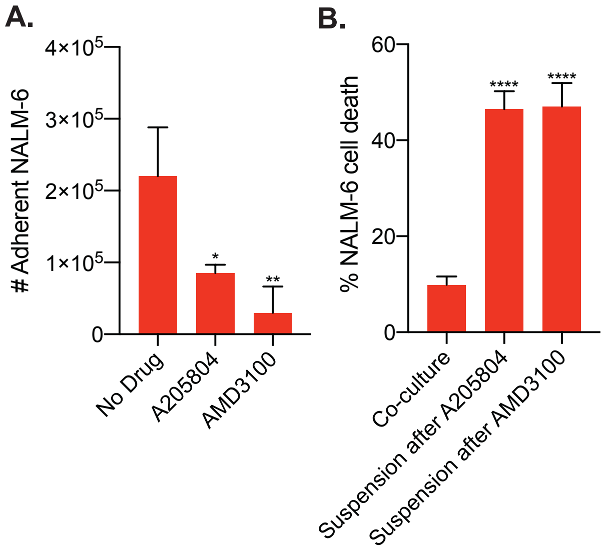
